# Impact of a 7-day homogeneous diet on interpersonal variation in human gut microbiomes and metabolomes

**DOI:** 10.1101/2021.10.29.465965

**Authors:** Leah Guthrie, Sean Paul Spencer, Dalia Perelman, Will Van Treuren, Shuo Han, Feiqiao Brian Yu, Erica D. Sonnenburg, Michael A. Fischbach, Timothy W. Meyer, Justin L. Sonnenburg

**Affiliations:** Department of Microbiology and Immunology, Stanford University School of Medicine, Stanford, CA, 94305, USA; Division of Gastroenterology and Hepatology, Stanford University, Stanford, CA, 94305, USA; Stanford Prevention Research Center, Department of Medicine, Stanford School of Medicine, Stanford, CA 94305, USA; ChEM-H, Stanford University, Stanford, CA, 94305, USA; Chan-Zuckerburg Biohub, San Francisco, CA, 94158, USA; Department of Bioengineering, Stanford University, Stanford, CA, 94305, USA; The Department of Medicine, VA Palo Alto Healthcare System, Palo Alto, CA, 94304, USA; Center for Human Microbiome Studies, Stanford, CA, 94305, USA

**Keywords:** Microbiome, uremic solutes, microbiome dependent metabolites, dietary intervention, precision medicine

## Abstract

Metabolism of dietary compounds by the gut microbiota generates a vast array of microbiome-dependent metabolites (MDMs), which are highly variable between individuals. The uremic MDMs (uMDMs) phenylacetylglutamine (PAG), *p*-cresol sulfate (PCS) and indoxyl sulfate (IS) accumulate during renal failure and are associated with poor outcomes. Targeted dietary interventions may reduce toxic MDMs generation; however, it is unclear if interindividual differences in diet or gut microbiome dominantly contribute to MDM variance. Here we use a 7-day homogeneous average American diet to standardize dietary precursor availability in 21 healthy individuals. Notably, the coefficient of variation in three uMDMs of interest, PAG, PCS, and IS (primary outcome), did not significantly decrease. The majority of circulating MDMs maintained variation despite identical diets. These results highlight the highly personalized profile of MDMs and limited contribution of short-term dietary heterogeneity, suggesting that dietary modification may need to be paired with microbial therapies to control MDM profiles.

## Introduction

Dietary change is known to impact the human gut microbiome and physiological status including influencing metabolic and immune parameters^1–3^. Human diets are sources of vast environmental exposure and are substrates for gut microbiota metabolism, which generates over 30,000 microbiome-dependent metabolites (MDMs)^4^. MDMs produced as a consequence of gut microbiota metabolism are associated with cardio-renal diseases^5,6^, obesity^7^, metabolic syndrome^8^ and cancer^9^, reach high concentrations (mM) in systemic circulation^10^, and most are cleared by the kidneys and eliminated in urine. When kidney function declines, MDMs accumulate as some of the most abundant uremic solutes (uMDMs)^10,11^, which have well defined mechanisms of toxicity^12–14^, and are difficult to clear via dialysis alone^10^. The levels of two uMDMs, p-cresol sulfate (PCS) and indoxyl sulfate (IS), are highly variable between individuals, but stable within an individual over time^15^, indicating precision strategies to reduce their level of production could be clinically beneficial. While targeted dietary interventions are an attractive approach to modulate the accumulation of potentially toxic MDMs, the extent that diet can be used to modulate and control MDM production remains incompletely understood.

Several interventions have investigated the influence of diet modifiable factors on MDM levels. Protein restriction was shown in early studies to ameliorate uremic symptoms and also to reduce urinary excretion of compounds including PCS, consistent with many uMDMs being amino acid-derived^16,17^. Increasing the intake of fiber, defined as carbohydrates that are not absorbed in the small intestine, has been hypothesized to program the microbiota to reduce degradation of amino acids^18,19^. Vegetarians produce less PCS and IS than omnivores, coincident with higher fiber and less protein intake^15^. In hemodialysis patients, a randomized trial has shown that daily intake of 18 g of fiber in the form of resistant starch for 6 weeks reduced plasma levels of IS^19^.

We carried out the Microbiome Individuality and Stability Over Time (MISO) study to investigate interpersonal variation in diet derived MDM levels when diet is homogenized and highly standardized. MISO study participants metabolomes and metagenomes were profiled before, during, and after undergoing a 7-day standardized homogeneous diet using habitual dietary data, metabolomic profiling of urine, feces and plasma and stool metagenomics. We designed the study to add a layer of stringency not previously achieved for microbiome focused studies, even in the most highly controlled in-patient diet intervention studies reported^3,20^. Even when participants are receiving the same meals as one another, meal-to-meal variation (e.g., different content of breakfast vs. lunch) combined with variation in gut motility can result in differences in the chemical environment experienced by two participant’s respective microbiomes at a given sampling time point. Therefore, interindividual differences in, for example, serum metabolites, are difficult to attribute to temporal diet variation vs. individualized aspects of the microbiota. Our study, in providing a uniform chopped salad format for 7 days ensures that bite-after-bite is homogenized between participants; therefore, it is expected that dietary chemical flow through the colon should be as uniform as possible between participants over the course of sampling during the intervention period. A reduction in interindividual variation in the urine levels of uMDMs, indoxyl sulphate (IS), p-cresol sulfate (PCS) and phenylacetylglutamine (PAG), uremic solutes that are efficiently excreted in the urine via the kidneys in healthy individuals and are associated with cardiovascular and renal toxicity, served as the primary outcome^21^.

As factors that shape intestinal microbiota composition and its metabolic output are incompletely understood, the homogenous dietary intervention helped to identify both dietary and host factors impacting microbiota composition and function. We identified host identity and age as dominant contributors to variability in microbiome-dependent metabolite levels broadly. Together, these findings demonstrate that microbiome-dependent solutes are variably controlled by direct and indirect combinatorial microbiome driven metabolic processes. Our approach identifies MDMs for which dietary strategies may be effective at reducing metabolite levels for clinical benefits and identifies MDMs for which the microbiome or other individualized factors play a dominant role over this dietary intervention. The results suggest microbiome reprogramming or alternative dietary strategies may be required for changing much of the metabolic output of an individual’s gut microbial community.

## Results

### Overview of study design and dietary intervention

We performed a controlled feeding study to quantify the impact of a homogeneous diet on interpersonal variability in the excreted levels of MDMs in a healthy cohort (ClinicalTrials.gov Identifier: NCT04740684). Of the 61 individuals assessed for eligibility, 26 were enrolled and 21 participants completed the study (**Figure 1A**). The final cohort were adults (age 48 ± 14 y [mean ± SD]), with a mean body surface area (BSA) of 1.93 ± 0.27 m2 (**Table S1**). We enrolled healthy volunteers with diverse habitual diets, to understand the effects of homogenizing their diet, regardless of their baseline diet. We quantified and compared MDM levels in enrolled participants during three phases: the baseline diet (BD) phase (days 1-14), the homogeneous diet (HD) phase (days 15 - 21) and the washout (WO) phase (days 22-28) (**Figure 1B**). During the 14-day BD phase, study subjects were instructed to maintain their habitual diet and food logs were used to assess their dietary patterns. Participants showed diverse habitual diets, defined by a coefficient of variation CV greater than 25 % of macronutrients (**Figure S1**). During the 7-day HD phase, study participants consumed single serve portions of a standardized diet *ad libitum* (**Table S2**) that was prepared in a commercial kitchen and packaged in 295 g portions.

**Figure 1.**
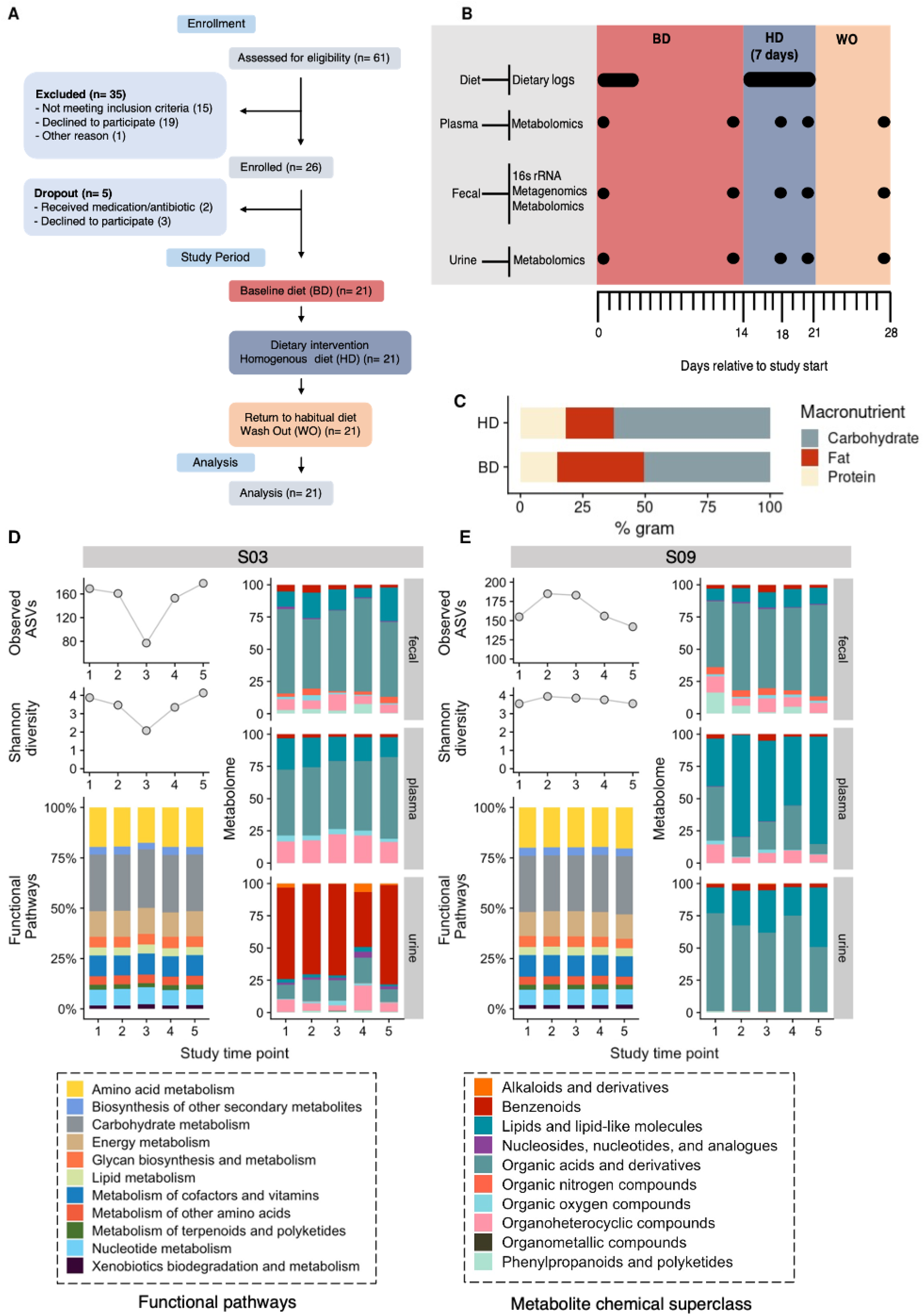
MISO study enrollment, design, and data collection. **(A)** CONSORT Flow diagram of participant enrollment and analysis in the MISO study. **(B)** The 4-week study overview timeline, sample types collected, timepoints of sample collection, and corresponding experimental platforms. **(C)** Average diet macronutrient composition during the baseline diet (BD) across the cohort and intervention homogenous diet (HD) study phases represented in percent grams. No baseline diet was available for participant 23. **(D)** Longitudinal data collected at study time points for MISO study participants S03 and S09 as illustrative examples. Observed ASVs and Shannon diversity were determined using 16S amplicon sequencing of fecal samples. Fecal metagenome functional profiles were derived from sequencing, mapping, and gene alignments. Urine and plasma metabolomes were quantified using a microbiome-focused metabolomics pipeline; metabolites were mapped to their structure-based chemical taxonomy (see Methods).

The HD was designed to recapitulate the diet quality, and specifically the fiber and macronutrient ranges^22^, common to adults in America based on the National Health and Nutrition Examination surveys and additional studies^22^. Although each participant consumed identical diets during HD (i.e., same composition and percentage of macronutrients), the amount of the HD consumed across participants varied as expected due to differences in individual caloric needs (we also did not want to confound the results with weight changes during this intervention) (**Figure S1)**. During the 7-day WO phase study subjects resumed their habitual diet. In addition to collecting detailed participant food logs during the BD and HD study phases, we collected stool, blood, and urine samples at five different timepoints, inclusive of all three study phases, for microbiome and metabolome profiling (**Figure 1B**). The overall macronutrient composition of dietary protein, carbohydrates and fats were relatively stable across the BD and HD study phases (37.5 vs 35.1% for fat, 17.0 vs 14.9% for protein and 45.8 vs 51.3% for carbohydrates), amounting to a modest decline in total protein between the BD and HD (**Figure 1C; Figure S1D**, paired Wilcoxon signed rank test, p = 0.03) and significant decrease in overall fiber during the HD phase (**Figure S1B**, paired Wilcoxon signed rank test, p < 0.0001).

We wished to define how interpersonal and intrapersonal (i.e., temporal) variability in microbiome-dependent metabolites (MDMs), genes and species were affected by the homogeneous diet. Therefore, we performed shotgun metagenomic sequencing and 16S rRNA sequencing on fecal samples and targeted metabolomics of uremic solutes on urine samples; and we used a novel microbiome-focused metabolomics pipeline that we recently reported^23^ on fecal, plasma and urine samples from each participant. The microbiome-dependent species, gene and metabolite profiles of two representative study participants demonstrates the limited changes in functional pathway relative abundance as well as the intrapersonal changes in observed species and alpha diversity that occurred during the HD intervention (**Figure 1D-E**). For the two highlighted participants, MDM-focused metabolomic profiling revealed extensive chemical diversity of the participants’ fecal, plasma and urine metabolomes, including notable intrapersonal stability of these profiles over the study (**Figure 1D-E**).

### A 7-day homogeneous diet does not reduce interpersonal variation in three amino-acid derived uMDMs

To characterize the impact of homogenizing diet over a 7-day period on interpersonal variation in the output of uremic microbiota-dependent metabolites (uMDMs), we profiled uMDM levels in urine using targeted mass spectrometry during the BD, HD and WO study phases. The primary outcome for this study was a 25% reduction of the coefficient of variation (CV) of 24 hr urinary excretion of at least 1 of 3 well-studied uMDMs that are derived from dietary amino acids: (*p*-cresol sulfate (PCS), indoxyl sulfate (IS), or phenylacetylglutamine (PAG)) measured and normalized to a body surface area of 1.73 m^2^ using the formula of Mosteller^24^, during the HD phase as compared to the BD phase (**Figure S2**). The first (BD phase) and fourth (HD phase) time points were compared for change in CV calculations. Urine IS levels decreased by 2.8 mg/day/1.73 with a CV decrease of 1.7% (p = 0.86). Urine PCS levels increased by 3.9 mg/day/1.73, with a CV increase of 11.2% (p = 0.20). PAG levels increased by 7.7 mg/day/1.73 and had a CV reduction of 0.05 % (p = 0.49). Therefore, although all three uMDMs exhibited changes in CV during the 7-day homogeneous diet, they did not uniformly decrease, and did not meet the primary endpoint as each of the 3 metabolites had less than a 25% reduction in interpersonal variation as measured by %CV (**Table 1**) and the % CV between time points 1 and 4 did not uniformly decline (**Figure 2A**).

**Table 1.** Coefficient of variation of uremic solutes

| Metabolite | Time point | Mean | SD | CV | *CV % |
| --- | --- | --- | --- | --- | --- |
| indoxyl sulfate | 1 | 23.8 | 11.9 | 0.49901 | -1.7% |
| indoxyl sulfate | 4 | 21.0 | 10.1 | 0.48164 |  |
| indoxyl sulfate | 5 | 28.5 | 15.4 | 0.54070 |  |
| p-cresol sulfate | 1 | 21.9 | 15.8 | 0.71993 | 11.2 % |
| p-cresol sulfate | 4 | 25.8 | 21.5 | 0.83168 |  |
| p-cresol sulfate | 5 | 29.6 | 20.2 | 0.68478 |  |
| phenylacetylglutamine | 1 | 84.0 | 55.0 | 0.65436 | -0.05 |
| phenylacetylglutamine | 4 | 91.7 | 59.9 | 0.65383 |  |
| phenylacetylglutamine | 5 | 107.0 | 74.7 | 0.69846 |  |
| Hippuric acid | 1 | 261.3 | 194.0 | 0.74231 | -27.9 % |
| Hippuric acid | 4 | 95.0 | 44.0 | 0.46304 |  |
| Hippuric acid | 5 | 277.4 | 212.6 | 0.76654 |  |

\* time point 1 vs time point 4

**Figure 2.**
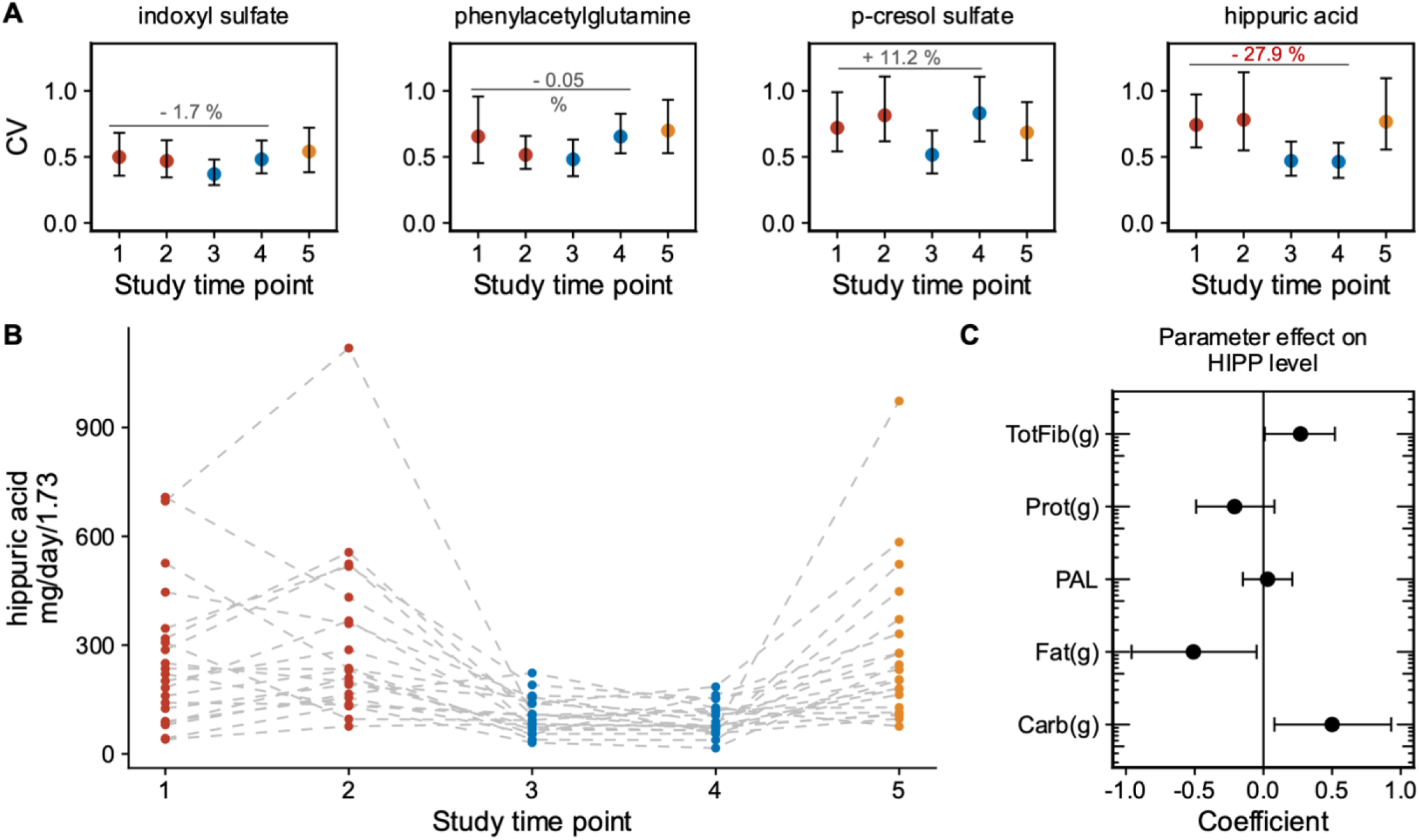
HD diet results in reduction of interpersonal variation in hippuric acid but not other uMDMs. **(A)** Coefficient of variation (CV) with 95 % confidence interval in urine uremic solutes indoxyl sulfate (IS), phenylacetylglutamine (PAG), *p*-cresol sulfate (PCS) and hippuric acid (HIPP) at study time points across the BD, HD, and WO phases. **(B)** Urine levels of hippuric acid (HIPP) determined by targeted LC-MS during the baseline diet (BD), homogenous diet (HD) and wash out (WO) (****p < 0.0001; paired Wilcoxon signed rank test). **(C)** Effect sizes of dietary macronutrients and microbial enzymes quantified by a mixed-effects model (along with a 95% CI) show that higher total fiber (p = 0.042) and carbohydrates (p = 0.036) supports higher levels of hippuric acid (HIPP) while phenylalanine ammonia-lyase (PAL) modestly improve HIPP levels, whereas total protein (p =0.03) detracts from HIPP levels (linear mixed-effect model on log10-transformed enzyme abundance data p = 0.04).

### Dietary fiber predicts urine hippuric acid levels

Notably, urine urea nitrogen, an indicator of total protein breakdown, decreased during the intervention phase (**Figure S1F**, paired Wilcoxon signed rank test, p = 0.01 with a CV reduction of 12.7%; p = 0.0018), consistent with the decreased protein intake by participants during HD (**Figure S1D**). These data indicate that the reduction in total protein was not sufficient to reduce interpersonal variance in these amino acid-derived metabolites. Given that we saw a reduction in total consumption of fiber during the intervention HD phase (**Figure S1B**, paired Wilcoxon signed rank test, p < 0.0001), we wondered whether other uMDMs derived from precursors found in fiber were sensitive to the HD phase of the study. Specifically, we analyzed levels of hippuric acid (**Figure 2B**), known to respond dynamically to dietary intervention^25,26^, and can be derived from both plant polysaccharides (e.g., fiber) and protein sources. During the HD phase, the mean levels of hippuric acid decreased and the overall interpersonal variation decreased by 27.9 % difference in hippuric acid CV between time point 1 (BD) and time point 4 (HD) (**Figure 2A**, p = 0.011). During the WO phase, the CV returned to baseline levels, indicating that dietary heterogeneity between participants is a key determinant in hippuric acid production and variation. We next hypothesized that the decrease in dietary fiber and intrinsic microbiome capacity shaped hippuric acid levels during the HD phase. Using a mixed-effects model that considered dietary macronutrient intake and microbiome genes as covariates and controlling for host parameters (i.e., biological sex, age, and body surface area), we quantified the role of diet and bacterial metabolism (metagenomic data) on the levels of hippuric acid. Total dietary fiber, carbohydrate levels, and the abundance of the microbiome encoded enzyme responsible for producing a precursor, phenylalanine ammonia-lyase (PAL), all positively contribute to hippuric acid levels based on the computed effect sizes (**Figure 2C**, linear model, p = 0.04). For IS, PCS and PAG there was no significant relationship between metabolite levels, diet, and microbiome-encoded abundance of genes involved in the respective metabolite’s generation (**Figure S2**). These findings indicate that the observed reduction in the CV and urine levels for hippuric acid is dietary precursor dependent, consistent with the decrease in variability observed between study participants during the HD phase.

### The HD intervention reduces inter- and intra-personal variation in microbiome composition, function and metabolic output

We leveraged the study design and participant samples to explore beyond the uMDMs and examine how a homogeneous diet impacts a broader array of microbiome-dependent metabolites, genes, and microbial composition. Metabolomic data for urine, plasma, and fecal samples resulted in 168, 183, and 320 different metabolites, respectively, observed in 80% of participants for at least one time point. We evaluated the % change in CV between BD phase time point 1 and HD phase time point 4 for all fecal, urine, and plasma metabolites present in ≥80% of a given sample type (see Methods). We identified 45 fecal, 15 plasma and 16 urine metabolites with ≥25% reduction in interpersonal variation (CV) during the HD phase (**Figure 3A**). Although some of the metabolites identified by this pipeline are independent of microbiota metabolism (e.g., derived from host, diet, or medication), known urine MDMs, *p*-cresol glucuronide and indole-3-acetic acid (IAA) are colon-derived uremic solutes^8^, that share precursors with PCS and IS respectively, and show a 66.4 % and 37.9 % reduction in CV in urine levels during HD, respectively; IAA also shows a reduction in fecal CV (27.6 %). While variation in overall microbiome composition, function and metabolic output were largely independent of study phase (**Figure S3**), this analysis suggests that unifying diet may reduce interpersonal variation for a subset of uremic solutes. To better understand the impact of the HD diet on all microbiome-dependent data types collected in the study, we calculated the Bray-Curtis dissimilarity (as a measure of uniqueness: the higher the value the more dissimilar from other samples) of participant microbiome composition (16S rRNA), fecal metagenomes, and metabolomes of urine, plasma and stool. Data were collected at all study time points, reflecting all three phases of the study. 16S rRNA amplicon sequencing resulted in an average of 11,422,190 (range 2419034 - 71084760) reads per sample, an average of 252 ASVs per sample (range 77-331), and 534 ASVs that occurred in 80% of participants for at least one time point (a criteria set for further analysis; see STAR methods). We found that interpersonal variation in microbiome composition and fecal metabolomes declined in the HD phase relative to the BD phase and increased during the WO phase relative to HD phase (**Figure 3B,C**; paired Wilcoxon signed rank test, p = 0.002 and p < 0.003; respectively). These data illustrate that a small but measurable amount of variation between people in the compositional features and functional output of the gut microbiome are driven by differences in diet.

**Figure 3.**
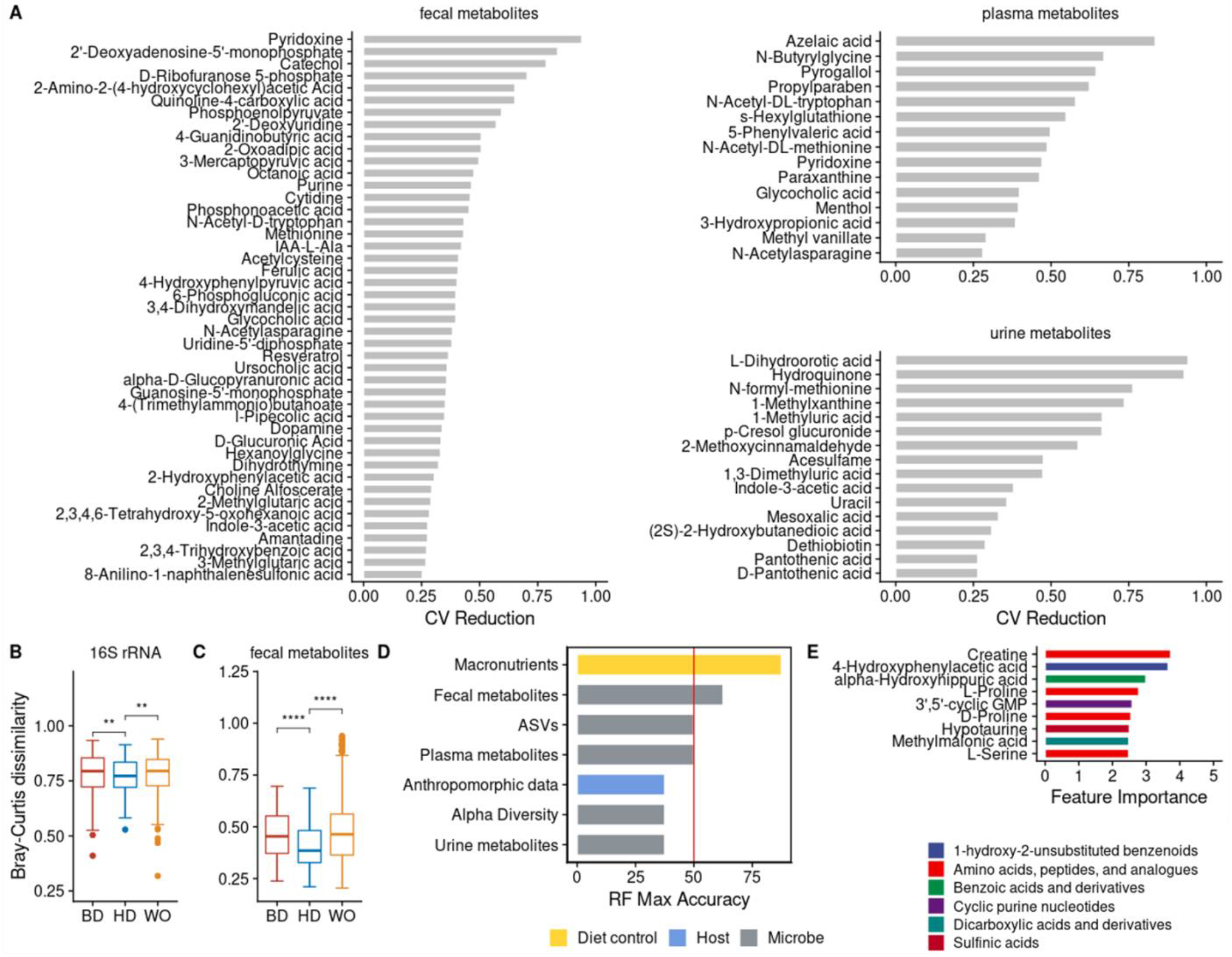
Interpersonal variation in microbiome composition and fecal metabolic output declines during HD intervention. (**A**) Fecal, plasma and urine metabolites with ≥25% reduction in CV across the cohort comparing timepoint 1 (BD) with timepoint 4 (HD). (**B**,**C**) Boxplots showing microbiome similarity measures for (**B**) ASV-level and (**C**) fecal metabolome based on Bray-Curtis dissimilarity metrics across study timepoints 1 (BD), 4 (HD) and 5 (WO). **(D)** Accuracy of leave-one-out cross-validation (LOOCV) of random forest models predicting study phase using different data types from study time points in BD (1 or 2) versus HD (3 or 4); diet macronutrient data (yellow), microbe-enriched data (grey) and anthropomorphic data from the host (blue). **(E)** Percent importance of individual fecal metabolites contributing to the model. Metabolites are colored by their Subclass level chemical ontology.

We generated random forest models to determine which microbiome-associated data types were useful in differentiating participant samples into BD vs HD phases. As a positive control, we developed a model based on participant macronutrient and fiber data, which at 87.5 % accuracy, had the highest accuracy for distinguishing BD (1 or 2) versus HD (3 or 4) study time points (**Figure 3D**). The best microbiome-associated data to classify participants by study phase were the fecal metabolomic profiles, with 62.5 % accuracy (**Figure 3D**). The fecal metabolite most explanatory for study phase was host derived creatine, followed by 4-hydroxyphenylacetic acid, an MDM of tyrosine (**Figure 3E**; time point 2 versus time point 3, LOOCV). Plasma and urine metabolomic profiles, and microbiome compositional features were no better than chance in classifying samples into the correct diet phase.

Previous studies have demonstrated a dominant role of diet and environment in dictating features within the microbiome^27,28^. We used permutational multivariate analysis of variance (PERMANOVA), to quantify the extent of variance in microbiome composition, function, and metabolic output explained by each host- and diet-associated variable that we measured. The majority of variation in microbiome features measured, which include species and genes found within the microbiome, and plasma, urine, and fecal metabolites, is due to host identity (**Figure 4A**), consistent with previous work demonstrating individuality of the microbiome in longitudinal dietary intervention studies^29–32^. Age (15%), body surface area (8%), and biological sex (3%) also explain a significant portion of variance across all data types (p < 0.05). Host identity has a greater impact on plasma (47%) and urine (45%) derived host-microbe metabolomes compared to participant fecal metabolomes (23%). Age contributes to 11% of the plasma host-microbe metabolome, but not to the urine or fecal metabolomes, consistent with previous reports of age-related plasma metabolomic profiles^29,33,34^. Overall, we found that the study time point explains 1-2% of the variance in microbiome composition, function, and metabolome.

**Figure 4.**
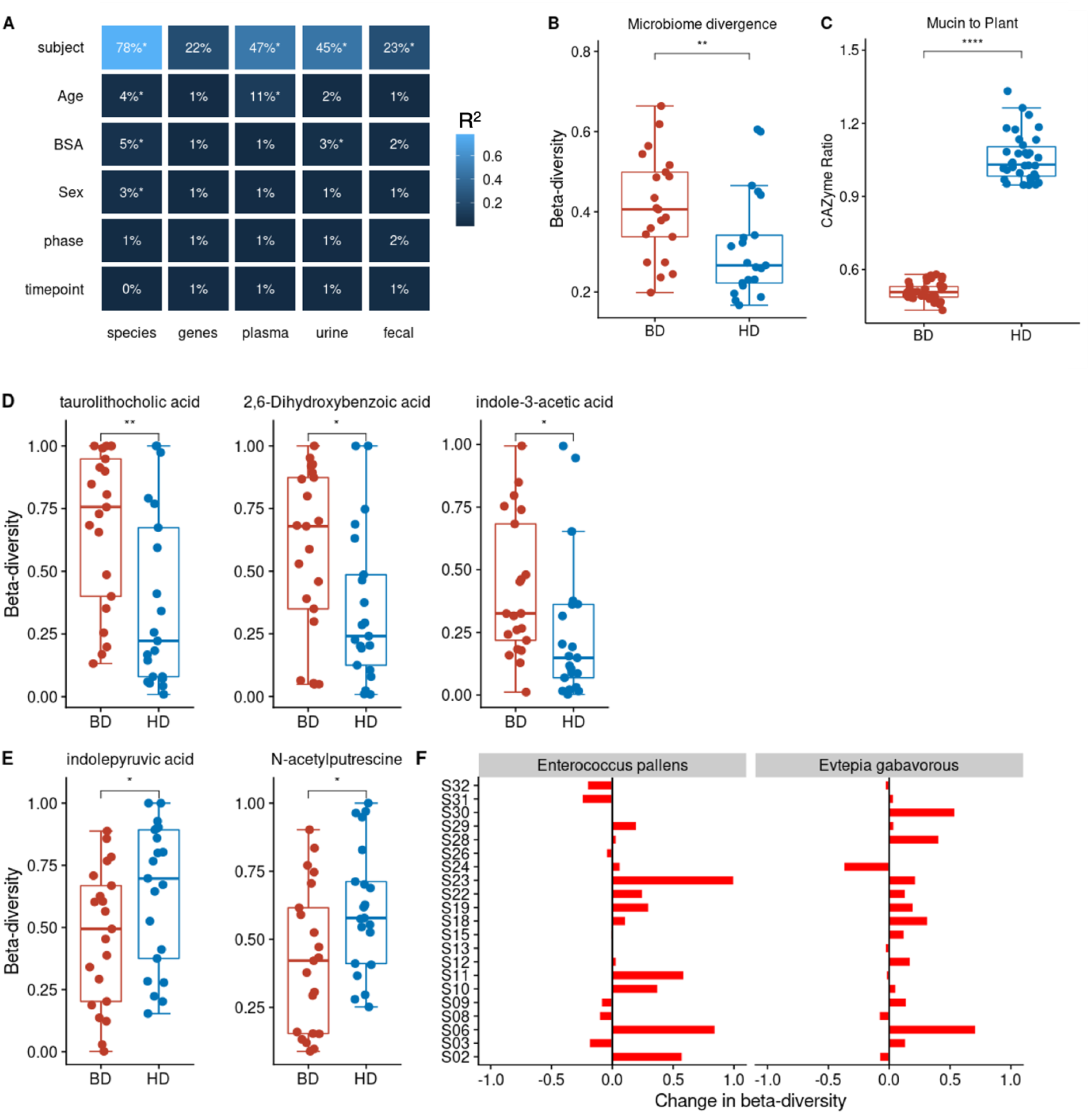
Intrapersonal variation in microbiome composition, function, and fecal metabolic output shifts after the HD intervention, but host identity explains most variance. **(A)** Host identity followed by age are the largest factors contributing to variation in microbiome composition, function and metabolomes in the MISO Study based on PERMANOVA (*p<0.05). Additional significant study variables shown capture < 10% of variation. Variance explained (R^2^) is also indicated by blue color scale. **(B)** Microbiome compositional intra-personal divergence collapses between the baseline (BD) and homogenous diet (HD) diet (paired Wilcoxon signed rank test,**p < 0.001;). **(C)** Microbiome mucin-to-plant carbohydrate active enzyme ratios (CAZyme) shifts between the BD and SD diet (paired Wilcoxon signed rank test,****p < 0.0001). **(D)** Levels of fecal metabolites, taurolithocholic acid, 2,6,-dihydroxybenzoic acid and indole-3-acetic acid levels are more similar during the HD (metabolite beta-diversity; Bray-Curtis distance) than the BD. Metabolite levels are based on 2 timepoints per study phase. **(E)** Fecal indole-3-pyruvic acid and N-acetyl putrescine increased in intrapersonal divergence during the HD phase (metabolite beta-diversity; Bray-Curtis distance). **(F)** Participant specific changes in beta-diversity for agmatinases from *Enterococcus pailens* (p < 0.05) and *Evtepia gabavorous* (p < 0.05) defined by the difference in beta-diversity between HD and BD phases. Specific microbial agmatinases involved in the production of N-acetyl putrescine increased in intrapersonal divergence during the HD phase (gene beta-diversity; Bray-Curtis distance) based on relative abundance counts of mapped agmatinase protein sequences against participant metagenomes.

We expected that if diet drives the decrease in interpersonal variability in some microbiome features between BD and HD phases, we would also observe that intrapersonal differences (i.e, temporal changes within study participants) would be greater in the BD phase, due to day-to-day variation in diet, in comparison to the HD phase. For each participant we calculated a Bray-Curtis dissimilarity matrix for the microbiome associated data types and computed the beta-diversity of each by comparing the difference in dissimilarity values for the two time points collected in the BD and HD phase. This approach allowed us to assess whether the homogeneous diet decreased overall interpersonal variation in these high-dimensional datasets. Participant microbiome 16S composition beta-diversity decreased significantly in the HD phase (**Figure 4B**, paired Wilcoxon signed rank test, p < 0.001). However, overall microbiota Shannon alpha diversity, a measure of species richness and diversity, did not decrease during the HD study phase or change cohort-wide over the study (**Figure S4**). These data are in keeping with the individuality and temporal stability of microbiome alpha and beta diversity of other short-term dietary interventions studies^28,35^.

Decreased dietary fiber consumption results in a shift in carbohydrate utilization within the microbiome towards mucin glycan degradation and away from plant carbohydrate degradation^30,36–38^. Therefore, we hypothesized that the decrease in fiber during the HD phase would be reflected in an increased ratio of genes predicted to encode mucin-relative to plant-glycan-degrading enzymes in participants’ metagenomes. We assigned glycoside hydrolases and polysaccharide lyases (i.e., carbohydrate active enzymes, or CAZymes) represented within participant metagenomes to mucin or plant carbohydrate degradation. An increase in the ratio of mucin-to-plant CAZYme genes in the HD relative to the BD study phases is apparent, consistent with a shift towards mucin consumption in response to the low fiber HD diet (**Figure 4C**, paired Wilcoxon signed rank test, p < 0.0001). Similar differences in microbiome-encoded carbohydrate genes have been found in studies of human diet with low fiber^38,39^.

To further characterize the relationship between the dietary intervention and MDMs, we next investigated intrapersonal divergence in participant fecal, urine and plasma metabolomes. Only three fecal metabolites decreased between the BD and HD phase, taurolithocholic acid, 2,6-Dihydroxybenzoic acid and indole-3-acetic acid (**Figure 4D**). Interestingly, two additional fecal metabolites, indole-3-pyruvic acid and N-acetyl putrescine, increased in intrapersonal variation despite identical dietary input during HD (**Figure 4E**), leading us to hypothesize that microbiome metabolism, rather than diet, plays a large role in their variation. Notably, all metabolites have consistent low levels of autocorrelation suggesting that the observed shifts were not due to time differences between sampling timepoints (**Figure S5**). To distinguish between the possibility that the increase was due to normal microbiome temporal dynamics or those induced by the shift in microbiome capacity on the HD, we investigated the intrapersonal divergence in the specific microbiome encoded genes responsible for production of these two metabolites. We mapped participant metagenomic profiles to a protein database of specific microbial genes encoding aromatic amino acid (AAA) aminotransaminases and agmatinases. These classes of enzymes include those that mediate the conversion of tryptophan into indole-3-pyruvic acid and arginine to putrescine, respectively (putrescine is converted into N-acetyl putrescine by host enzymes). Agmatinases that map to *Enterococcus paliens* and *Evtepia gabavorous* increase in overall intrapersonal divergence in the HD phase relative to the BD phase (**Figure 4F**) consistent with increased within-person variation in N-acetyl putrescine during HD. No similar pattern for aminotransaminase divergence was detected. Combined, these results suggest that homogenizing dietary input across healthy adults for 7 days contributes to some changes in microbiome composition, function, and specific MDMs, however, is insufficient to significantly reduce interpersonal and intrapersonal variation in many other facets of microbiome composition function and metabolic output.

## Discussion

There is great interest in optimizing the production of microbiome-dependent metabolites relevant to human health, including minimizing those that contribute to uremic illness. For instance, microbiota generated products, *p*-cresol sulfate and indoxyl sulfate, derived from amino acids promote the progression and exacerbation of cardio-renal diseases^14,40,41^. The use of diet modification to alter specific aspects of microbiome functional output has shown some promise in previous studies^42–45^, but the extent to which inter-individual variation in MDM production is caused by a person’s microbiome vs. their habitual diet remains an open question. In this study, we employed a 7-day homogeneous diet (HD) modeled on an average diet in the U.S. to eliminate dietary variability between individuals while attempting to minimize perturbation when participants shifted to this dietary intervention. The homogeneous diet significantly reduced interpersonal variation in hippuric acid, consistent with the low amount of dietary fiber precursor, but had no significant effect on other well studied uremic solutes including PAG, IS and PCS. The inability of HD to decrease the cohort-wide variability in these three uMDMs by 25% (the primary endpoint) provides strong evidence that diet-independent individualized features of the microbiome and host play a critical role in dictating individual-specific levels of these metabolites.

We generated microbiome-focused metabolomics data and sequencing data to investigate how the diet intervention affects a broader array of MDMs, as well as microbiome composition and function. These analyses broaden our understanding of MDMs that are sensitive and insensitive to dietary variation. Interestingly, we found that the diet phase of the study contributed to less than 2% in the overall variance found in participant fecal metabolomes. Host identity and age were the dominant contributors to interpersonal variability. Our results support precision approaches to modify MDM levels and identify a small group of metabolites for which dietary strategies may be effective at reducing metabolite levels for clinical benefits. Notably 16S-based composition analysis and fecal metabolite profiles show a small but significant decrease in interpersonal variation during HD consistent with diet contributing to some measurable extent of individuality.

The design of our study allowed us to dissect the relative contributions of diet (i.e., macronutrients) and microbiome functional potential (i.e., encoded genes) towards the interpersonal and intrapersonal variation in microbiome features (i.e., microbiome-dependent uremic solutes, 16S-based composition, metagenomic functions, and metabolome profiles). We found that the modest reduction in protein between the BD and HD phase (average 82 grams per day for BD, and 67 grams per day for HD) did not correspond to a reduction in the protein- derived metabolites and uremic solutes, PCS, IS and PAG. Previous studies quantifying IS and PCS, found that a very low protein diet (20 grams per day) was necessary to reduce their levels^46–48^. We also found that tyrosine, phenylalanine, and tryptophan, which are precursors for p-cresol, phenylacetylglutamine and indoxyl sulfate; respectively, were stable across the BD and HD phase.

Our analysis reveals that a homogenous diet reduces interpersonal variation in a small subset of metabolites and has modest but detectable effects on microbiome community composition. This suggests that the short-term homogenizing of diet has a limited capacity to normalize individualized features of the microbiome. Previously, studies of the gut microbiome in healthy cohorts have found that microbiome-food relationships are highly personalized^35^. Controlled feeding studies, which allow for the precise accounting of diet composition, can produce more reliable diet-microbiome interactions and associations^3,20,49^; however, to date, controlled feeding studies have not specifically addressed the impact of homogenizing diet composition on interpersonal variation in microbiome features. Our data adds a stringent test to demonstrate that participants’ individualized responses to dietary interventions are likely governed in part by the individualized aspects of microbiome composition and function that are fairly recalcitrant to short-term change. A greater understanding of a participant’s habitual diet is critical to understanding how diet interacts with microbiome functionality to determine metabolite production, and the extent to which metabolite levels may respond to an intervention.

How changes in the chemical composition of diet can alter microbiome metabolite output needs to be tested under a variety of conditions (including extreme dietary control) and provision of specific chemical precursors) with an emphasis on understanding participants’ baseline dietary habits and microbiome functional capacity (metagenomes). Furthermore, future studies should focus on precision manipulation of the microbiome (e.g., microbial therapies) in combination with dietary interventions. To develop targeted approaches to modulate metabolite levels, factors contributing to individual MDM level variability should be understood. Thus, a detailed understanding of the relationship between dietary input and microbiota metabolism in shaping interpersonal variability provides a roadmap to develop diverse therapeutic approaches. Importantly, unifying how dietary intervention studies account for variation in participant habitual diets, record and characterize diet data, lifestyle factors, host genetics, as well as the range of metabolites measured^50,51^, will help fill gaps in understanding the utility of diet to modulate the production of specific solutes.

## Acknowledgements

We wish to thank the participants for their engagement and effort to enable this study. Chef Laura Stec developed the recipe given many constraints, organized and led the cooking and packaging sessions. This work was funded through philanthropic support and the NIH/NIDDK 2R01DK101674 (J.L.S, M.A.F., T.W.M) and R01 3R01DK08502510 (J.L.S) and L.G. was supported by NIH Training Grant T32AI007328 and the HHMI Hanna Gray Fellowship. S.P.S. was supported by the A.P. Giannini Foundation Research Fellowship and Leadership Award and the NIH Training Grant T32DK007056. S.H. was supported by Stanford Dean’s Postdoctoral Fellowship and NRSA F32AG062119. J.L.S and M.A.F are Chan-Zuckerberg Biohub investigators.

## Author Contributions

L.G., S.P.S., E.D.S and J.L.S., T.M., and M.F.A conceived of the study. L.G. and S.P.S performed data analyses. D.P. provided patient interaction, dietary counseling, and analysis. F.B.Y. and S.P.S led sequencing. W.V.T. and S.H. designed the mass spectrometry pipeline. L.G. conducted mass spectrometry analysis. T.M., D.P., S.P.S. and J.L.S. oversaw patient recruitment and study design implementation. L.G., S.P.S., E.D.S and J.L.S wrote the manuscript. All authors contributed to the critical review of the manuscript.

## Materials and Methods

### Recruitment and selection of participants

Participants were recruited from the local community through online advertisement in different community groups as well as emails to past research participants that consented to being contacted for future studies. The current study assessed 61 participants for eligibility. They completed an online screening questionnaire and a clinic visit between September 2018 and January 2019. The primary inclusion criteria included age ≥ 18 y and general good health. Participants were excluded if they had a history of active uncontrolled inflammatory bowel disease (IBD) including ulcerative colitis, Crohn’s disease, or indeterminate colitis, irritable bowel syndrome (IBS) (moderate or severe), infectious gastroenteritis, colitis or gastritis, *Clostridium difficile* infection (recurrent) or *Helicobacter pylori* infection (untreated), malabsorptive intestinal disease (such as celiac disease), major surgery of the GI tract, with the exception of cholecystectomy and appendectomy, in the past five years, or any major bowel resection at any time. Other exclusion criteria included a BMI ≥ 40, diabetes, renal disease, significant liver enzyme abnormality, pregnancy or lactation, smoking, a history of CVD, inflammatory disease, or malignant neoplasm. CONSORT clinical trial flow diagram of participant recruitment shown in Figure 1A and demographics table shown in Table S1. 21 participants (11 female sex and gender identifying, 10 male sex and gender identifying) were used for full analysis with an average age of 48 +/-14 years. All study participants provided written informed consent. The study was designed as an exploratory approach toward discovery of changes in the microbiota and the metabolome in response to a dietary intervention. The study was approved annually by the Stanford University Human Subjects Committee. Trial was registered at ClinicalTrials.gov, identifier: NCT04740684.

### Specimen collection

Stool was collected at five study time points over a four-week period of the study and kept in participants’ home freezers (-20°C) wrapped in ice packs until they were transferred on ice to the research laboratory and stored at -80°C. 24 hr urine collection was obtained on Day 0 and 13 (while on BD), Day 17 (while on HD), and Day 28 (during WO phase). Plasma and spot urine samples were obtained during research clinic visits throughout the study at indicated time points.

## METHOD DETAILS

### Intervention

The study period lasted 28 days and consisted of a 14-day period while participants consumed their habitual baseline diet (BD), followed by 7 days of provided homogenous diet (HD), and concluding with a 7-day washout period (WO) of return to their prior habitual dietary pattern. They were asked to keep detailed food logs for the first 3 days upon initiation of the study. During the HD phase of study, participants were asked to record the quantity of packets consumed daily.

### Formulation of Homogenous Diet (HD)

The Homogenous Diet (HD) was provided for days 15-21 for the study period (7 days total). It was designed to recapitulate the diet quality, and specifically the fiber and macronutrient ranges^22^, common to adults in America based on the National Health and Nutrition Examination surveys and additional studies^22^. The food provided was a nutritionally adequate diet, designed by a registered dietitian. It was prepared by a professional chef in a commercial kitchen by mixing and cooking foods purchased from grocery stores. The food mixture was divided into 295 g portions that included a 210 g portion of the homogenous diet in addition to 59 g Orange Juice and 26 g of cookies to be consumed with each meal.

### Distribution of Food and Monitoring of Consumption

Over the 7-day homogenous diet period, subjects were allowed to eat as much of the homogeneous diet as they wanted to meet their caloric needs. Subjects were asked not to eat anything other than the provided HD (no candy, snacks, etc.) and not to drink anything except water (no coffee, tea, sodas, or alcoholic drinks, etc.). The ingestion of the HD study diet in each individual subject terminated with the urine, stool, and blood collection on the seventh day of eating the diet. After termination of the HD study diet, participants returned to their habitual diet.

### Measurement of uMDMs

Solute excretion and nutrient consumption rates were corrected for body surface area calculated using the Mosteller formula^24^.

### 16S amplicon sequencing

DNA was extracted from stool using the DNeasy PowerSoil HTP 96 kit according to the manufacturer protocol and amplified at the V4 region of the 16S ribosomal RNA (rRNA) subunit gene and 250 nucleotides (nt) Illumina sequencing reads were generated. 16S rRNA gene amplicon sequencing data from stool samples were demultiplexed using the QIIME pipeline^52^ version 1.8. Amplicon sequence variants (ASVs) were identified with a learned sequencing error correction model (DADA2 method)^53^, using the dada2 package in R. ASVs were assigned taxonomy using the GreenGenes database (version 13.8). ɑ-diversity was quantified as the number of observed ASVs, Shannon diversity, or PD whole tree, in a rarefied sample using the phyloseq package in R (version 3.4.0). 16S enumeration data resulted in an average of 11,422,190 (range 2419034 - 71084760) reads per sample, average 252 ASVs per sample (range 77-331), and 534 ASVs that occurred in 80% of participants for at least one time point that were used for further analysis.

### Metagenomic Sequencing

DNA extraction for shotgun metagenome sequencing was done using the DNeasy PowerSoil HTP 96 kit as described in the 16S amplicon sequencing methods. For library preparation, the Nextera Flex kit was used with a minimum of 10ng of DNA as input and 6 or 8 PCR cycles depending on input concentration. A 12 base pair dual-unique-indexed barcode (CZ Biohub) was added to each sample and libraries concentration and size were quantified using an Agilent Fragment Analyzer. They were further size selected using AMPure XP beads (Beckman) targeted at a fragment length of 450bp (350bp size insert). DNA paired-end sequencing (2×146bp) was performed on a NovaSeq 6000 using S4 flow cells (CZ Biohub). The average target depth for each sample was 11.4 million paired-end reads. Data quality analysis was performed by demultiplexing raw sequencing reads and concatenating data for samples that required multiple sequencing runs for target depth before further analysis.

BBtools suite (https://sourceforge.net/projects/bbmap/)) was used to process raw reads and mapped against the human genome (hg19) after trimming, with masks over regions broadly conserved in eukaryotes (http://seqanswers.com/forums/showthread.php?t=42552). Exact duplicate reads (subs=0) were marked using clumpify and adapters and low-quality bases were trimmed using bbduk (trimq=16, minlen=55).

### Plasma, Urine and Stool metabolomics

Metabolites from three sample types (plasma, urine, and stool) metabolites were profiled using three complementary liquid chromatography-tandem mass spectrometry (LC–MS) methods designed to measure a broad range of microbial and microbe-host co-metabolites^23^. For all three sample types, samples were incubated for 5 minutes at room temperature and centrifugation at 5,000 × g for 10 minutes. Sample supernatants were then transferred, evaporated, and reconstituted in an internal standard mix in (50% Methanol). A Hydrophilic Interaction Liquid Chromatography (HILIC) method was used for the analysis of water-soluble polar metabolites in positive (HILIC-pos) ion mode, and two C18 column chromatography methods for measuring metabolites of intermediate polarity, in positive (C18-pos) or negative (C18-neg) ion mode. Raw data were processed, and compounds were annotated using MSDIAL software 3.988^54^. Analyses were conducted using the data obtained from all three LC-MS methods after removal of features observed in <80% of the samples and imputing missing values with half of the minimum observed measurement for each feature.

### Quantification of Alpha and Beta Diversity Measures

Alpha and beta-diversity measures were calculated using the phyloseq package^55^ in R. Alpha diversity was calculated at the ASV level. For ASVs, metagenomes, plasma metabolomes, urine metabolomes and stool metabolomes, Bray-Curtis dissimilarity was computed at each timepoint and for each subject using the vegdist function in the R vegan package^56^.

### Microbiome Functional Profiling

The assembled metagenomes were mapped against the following functional databases using USEARCH version 8 with an e-value cutoff of 1e-40 in order to ensure longer sequence hits for improved taxonomic and functional resolution: KEGG EC/KO (*n* = 2,000,708)^57^, Carbohydrate active enzyme (CAZy) (*n* = 7215)^58^, and a curated database of both cultured and metagenomically identified agmatinases and aminotransaminases developed in-house. The abundance of KEGG orthologous groups and modules were determined using the HUMAnN pipeline^59^ with default parameters.

### Metabolite Chemical Class Mapping

All metabolites were mapped to their superclass, class and subclass using the ClassyFire automated chemical classification based on compound InChIKeys60.

### Statistical Analysis

All statistical analyses were carried out in R. The coefficient of variation and 95% confidence interval for each uremic solute (IS, PCS, PAG and HIPP), at each study time point was determined using the R cvcqv package. Statistical difference in CV or beta-diversity between time points 1 and 4 and mean metabolite abundances between the BD and HD phases were determined using a paired Wilcoxon signed rank test. The autocorrelation decay was determined for each metabolite with significant changes in beta-diversity using the autocorrelation function in the timeseries R package.

### Linear Mixed Effects Modeling

To study the effect of specific dietary parameters and the microbiome on the levels of hippuric acid during the HD phase, when diet is homogenized, we used a linear mixed effects model of hippuric acids levels, modeled as a function of the macronutrients, fiber as well as microbiome encoded genes with time point specific random effects. Terms for biological sex, age and body surface area were included in the model as covariates.

### Recursive Feature Random Forest

Random forest regression models were built of the default set of 1000 trees, with the caret R package^61^ to predict the study phase based on microbiome features. Training was achieved through 10-fold cross validation with ASV data, fecal metabolome, plasma metabolome, urine metabolome, dietary data, and anthropometry data. The feature selection was performed by using the recursive feature elimination algorithm of the caret R package^55^. For the random forest model based on ASVs, 534 ASVs that occurred in 80% of participants for at least one time point that were used for further analysis. 320 fecal metabolites, 183 plasma metabolites and 168 urine metabolites were used to build separate models. The importance scores of features were determined based on the increase of prediction error when that feature was randomly permuted while all others were remained unchanged.

### PERMANOVA

To calculate the variance explained by each of our collected study factors we performed an Adonis test implemented in the R vegan package (v2.5.7 using adonis2) using 999 permutations, with random permutations constrained by using the “strata” option. We used Bray-Curtis dissimilarity as the distance measure for all five data types independently, which includes participants ASVs, metagenomes, plasma metabolomes, urine metabolomes and stool metabolomes.

## Main table titles and legends

Table 1. Primary and exploratory clinical trial outcomes.

Coefficient of variation at study time points 1 (BD phase), 4 (end of HD phase) and 5 (WO phase). A percent reduction in CV between time points 1 and 4 of 0.25 or greater meets the endpoint.

## Supplemental table titles and legends

Table S1, Related to Figure 1: Participant characteristics

Table S2, Related to Figure 1: Food composition of the homogenous diet intervention

## Supplemental figure titles and legends

**Figure S1.**
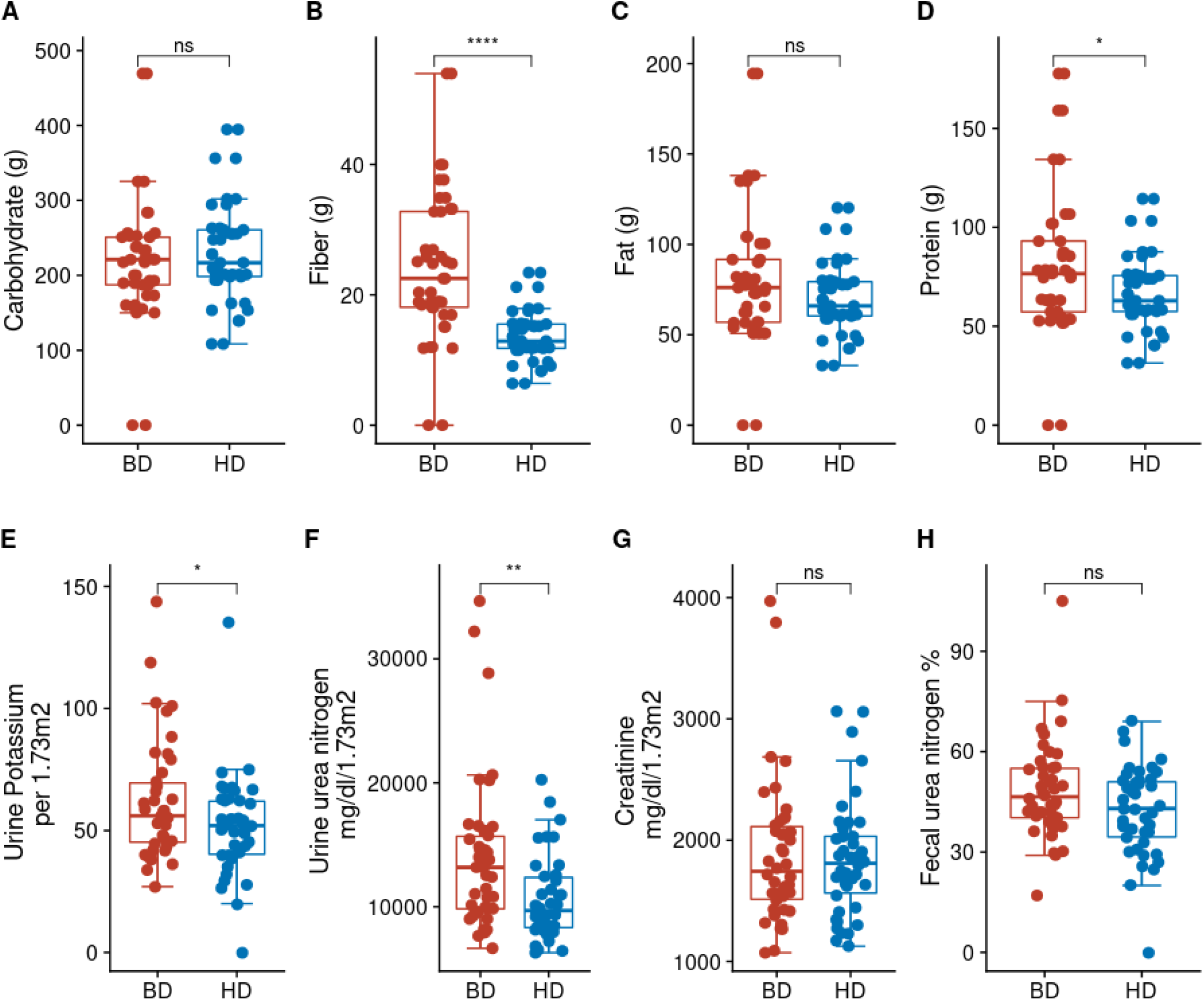
Related to Figure 1: Macronutrient and clinical parameters between BD and HD. Mean **(A)** carbohydrate, **(B)** fiber, **(C)** fat, and **(D)** protein intake during the BD and HD phases. Participants **(E)** urine potassium, **(F)** urine urea nitrogen, **(G)** creatinine, and **(H)** fecal urea nitrogen quantified at all study phases.

**Figure S2.**
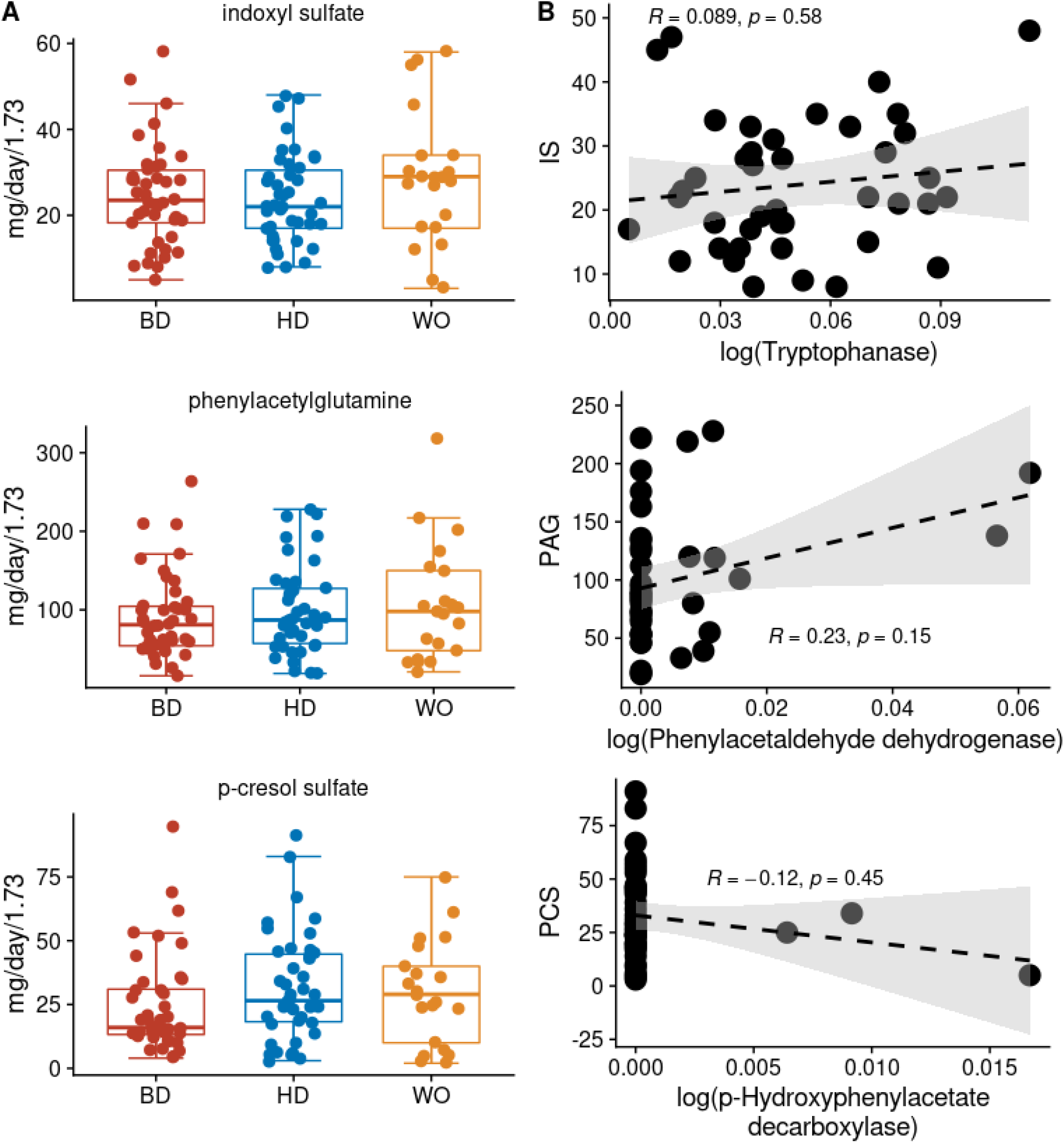
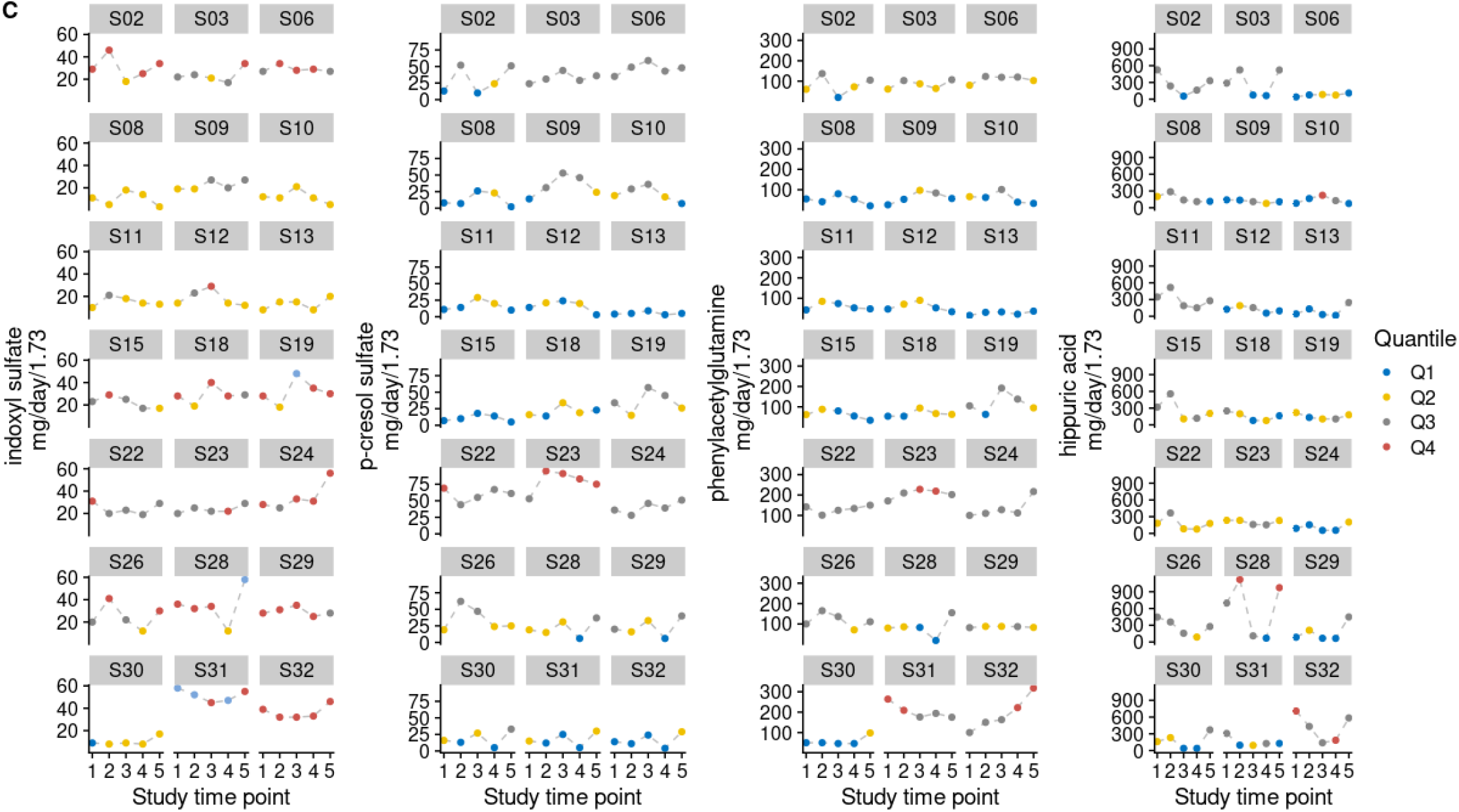
Related to Figure 2: Primary outcome uremic solute levels and correspondence with microbiome encoded metabolism. **(A)** Mean indoxyl sulfate (IS), phenylacetylglutamine (PAG), *p*-cresol sulfate (PCS) at the BD, HD, and WO phases. **(B)** Correlation between urine IS, PAG, PCS levels with the relative abundances of the microbiome-encoded enzyme involved in their production. **(C)** Uremic solute levels over the course of the MISO study by subject and colored by tertile.

**Figure S3.**
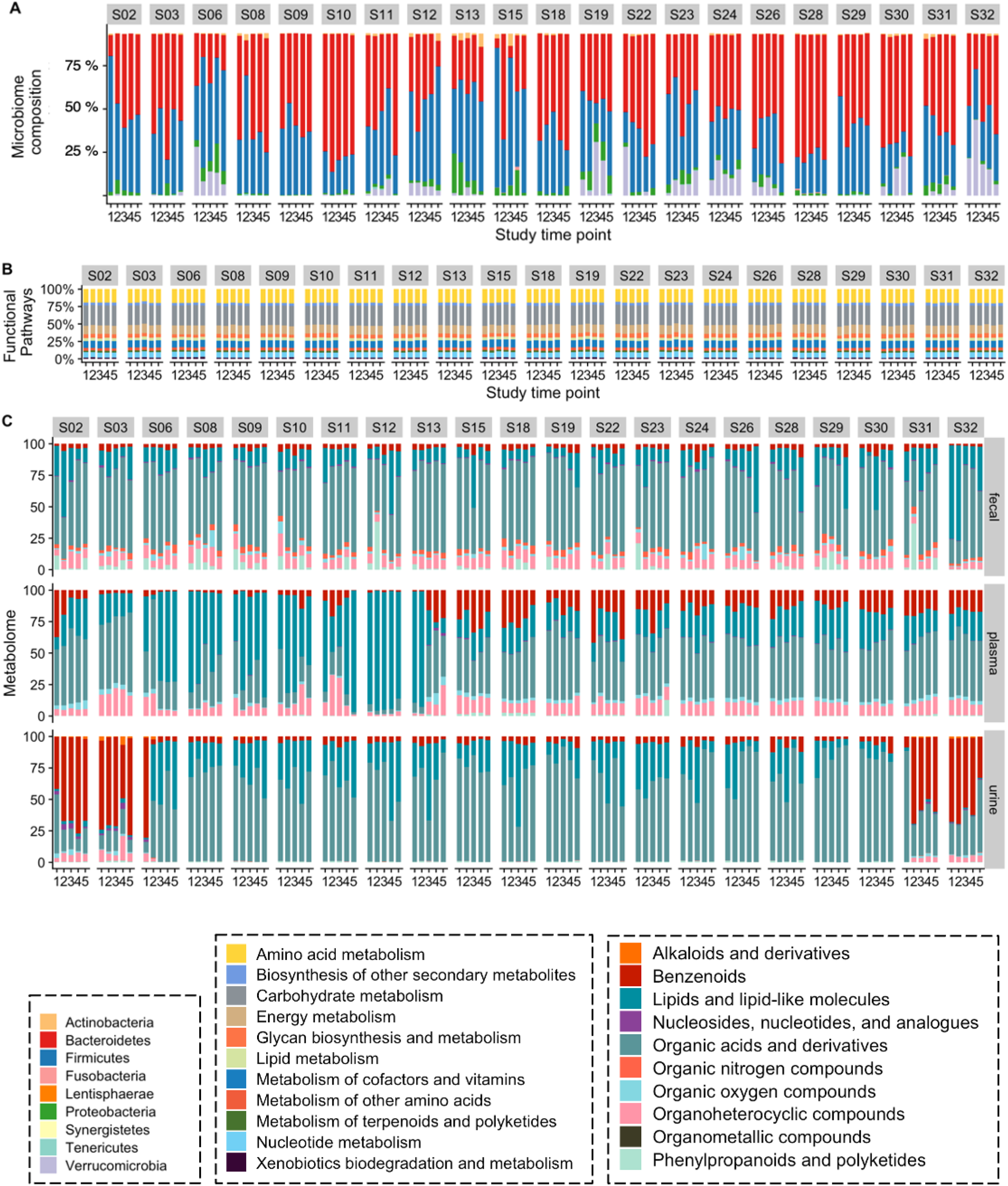
Related to Figure 3: Taxonomic, metabolic pathway, and metabolomes profiles for individual participants in the MISO study. **(A-C)** Longitudinal microbiome (**A**) composition (phylum level), (**B**) functional category relative abundance, and(**C**) fecal, plasma and urine metabolomes categorized by chemical class for MISO study participants.

**Figure S4.**
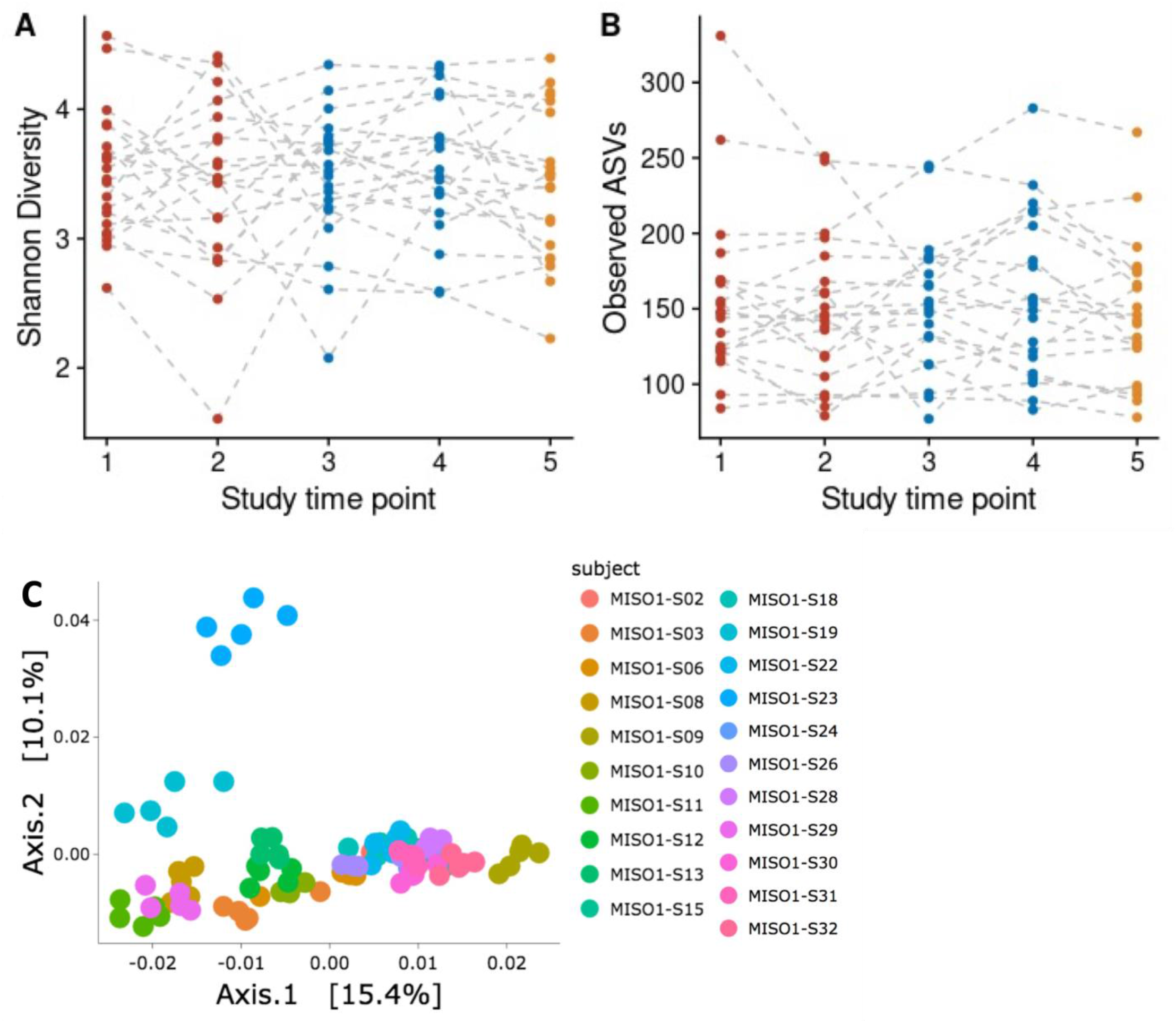
Related to Figure 4: MISO participants’ (A) Shannon Diversity, (B) ASV count and (C) personalized temporal variation in microbiome composition.

**Figure S5.**
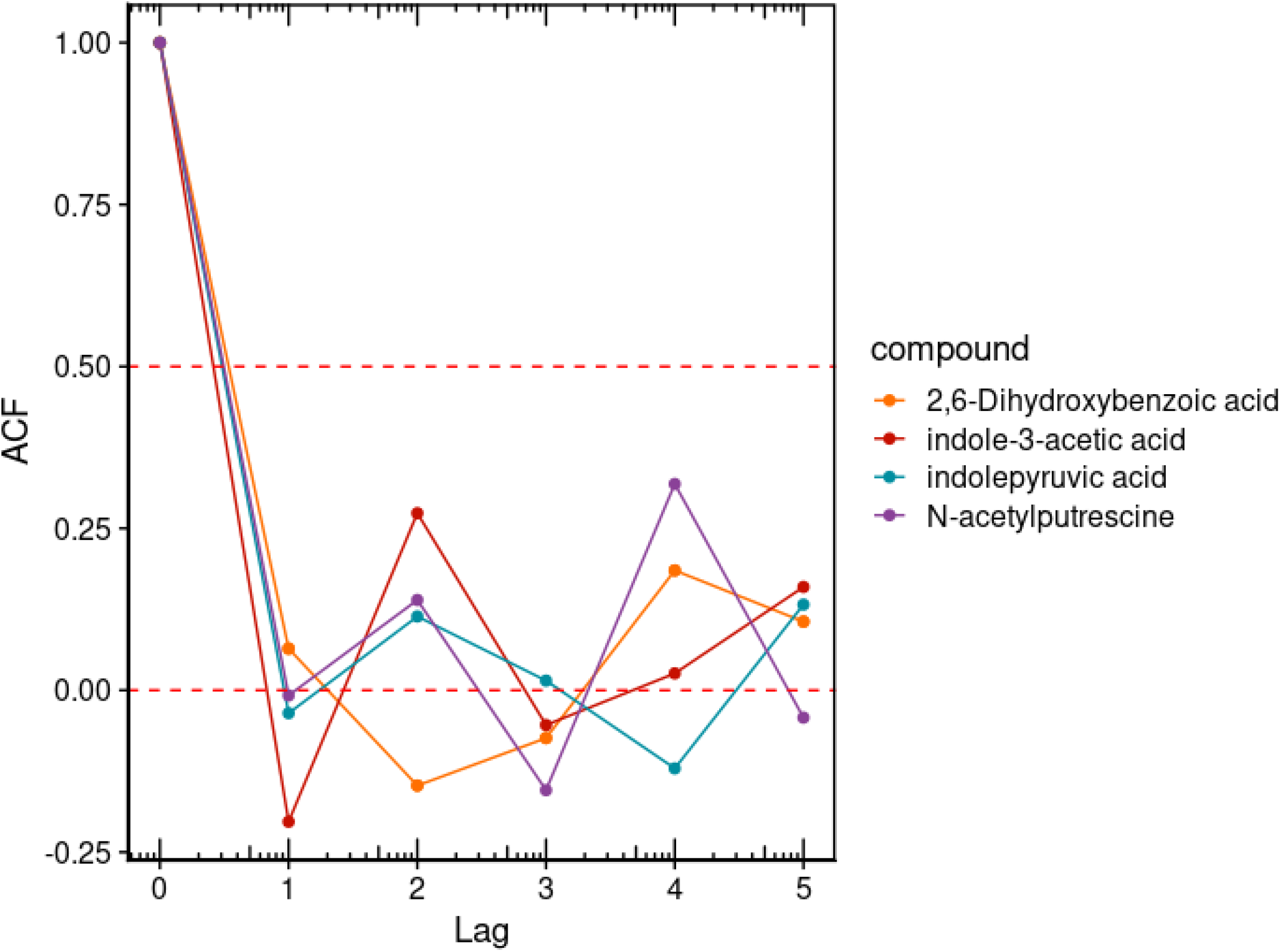
Related to Figure 4: Autocorrelation decay table for microbiome metabolites.

## Supplemental Tables

**Table S1.** Participant characteristics

| **subject** | **Sex** | **Age** | **BSA** |
| --- | --- | --- | --- |
| S02 | F | 56 | 1.9 |
| S03 | F | 55 | 2 |
| S06 | M | 28 | 2.1 |
| S08 | F | 27 | 1.8 |
| S09 | M | 23 | 1.7 |
| S10 | M | 46 | 1.8 |
| S11 | F | 58 | 1.6 |
| S12 | F | 27 | 1.7 |
| S13 | M | 34 | 2.8 |
| S15 | F | 30 | 1.9 |
| S18 | M | 46 | 1.9 |
| S19 | M | 50 | 2 |
| S22 | F | 54 | 1.7 |
| S23 | F | 53 | 1.7 |
| S24 | F | 54 | 2 |
| S26 | F | 62 | 1.8 |
| S28 | F | 45 | 1.8 |
| S29 | M | 58 | 1.7 |
| S30 | M | 67 | 2.1 |
| S31 | M | 75 | 2.3 |
| S32 | M | 54 | 2.4 |

**Table S2.**
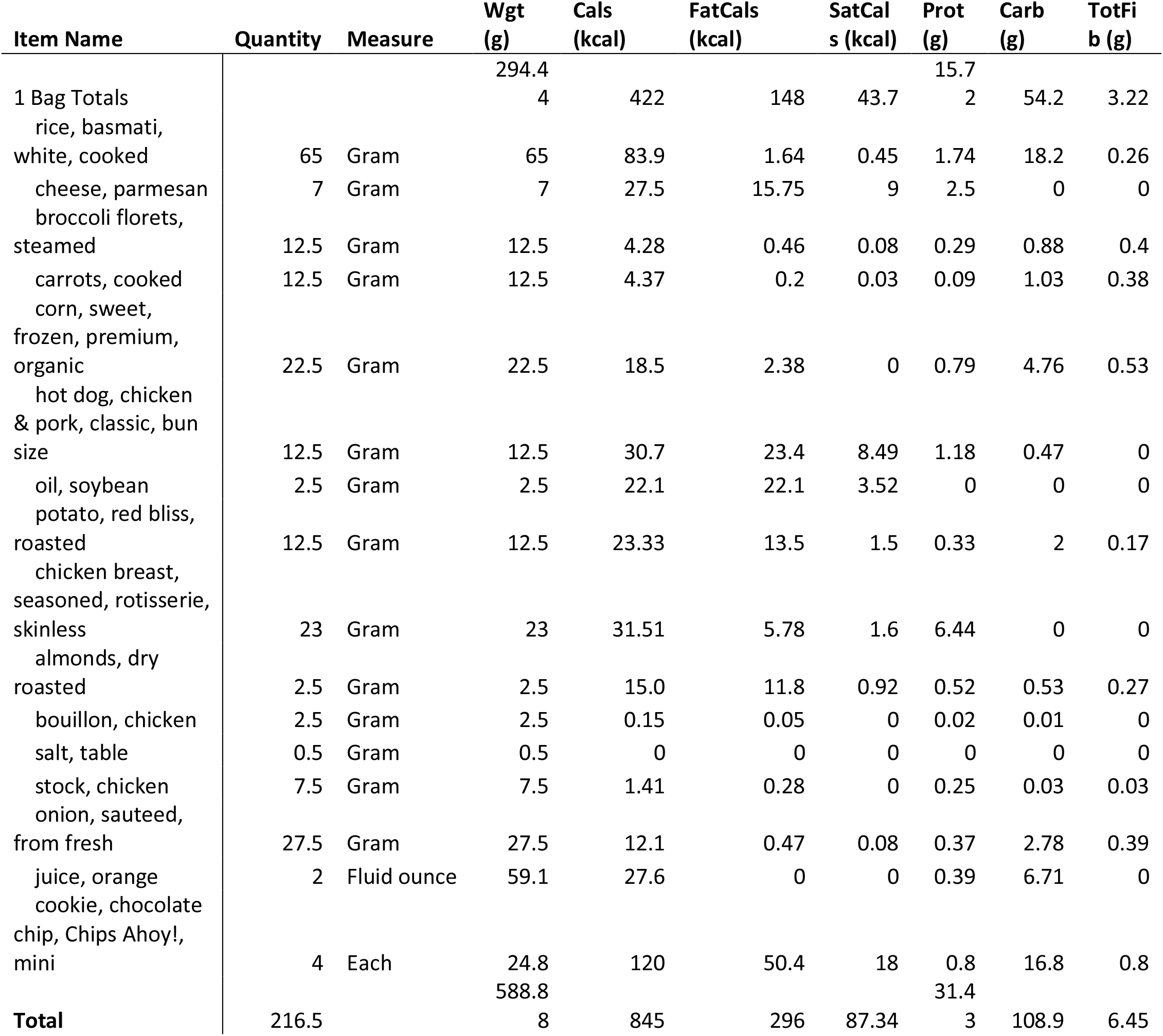
Food composition of the homogenous diet intervention

